# Olfactory receptor-dependent receptor repression in *Drosophila*

**DOI:** 10.1101/2021.04.27.441568

**Authors:** Kaan Mika, Steeve Cruchet, Phing Chian Chai, Lucia L. Prieto-Godino, Thomas O. Auer, Sylvain Pradervand, Richard Benton

## Abstract

In olfactory systems across phyla, most sensory neurons transcribe a single olfactory receptor gene selected from a large genomic repertoire. We describe novel receptor gene-dependent mechanisms that ensure singular expression of receptors encoded by a tandem gene array in *Drosophila*. Transcription from upstream genes in the cluster runs through the coding region of downstream loci to inhibit their expression in *cis*, via transcriptional interference. Moreover, one receptor blocks expression of other receptor proteins in *trans* through a post-transcriptional mechanism. These repression mechanisms operate in endogenous neurons to ensure their unique expression. Our data provide evidence for inter-olfactory receptor regulation in invertebrates, and highlight unprecedented, but potentially widespread, mechanisms for ensuring exclusive expression of chemosensory receptors, and other protein families, encoded by tandemly-arranged genes.

## Main text

The selective neuronal expression of olfactory receptors is fundamental to the sensory representation of odors in the brain. All current models explaining singular receptor expression in vertebrates and invertebrates invoke the binding of transcriptional activators to one receptor locus while other receptor genes are silenced through transcriptional repressors or repressive chromatin structure (*1–3*). Invertebrates are believed to rely only on deterministic, transcriptional codes to achieve specificity of receptor expression (*3–5*). By contrast, mammalian olfactory receptor regulation incorporates a feedback pathway, in which an expressed receptor protein prevents activation of other receptor genes via induction of the unfolded protein response to lead to stabilization of receptor gene choice and prevention of de-silencing of other receptor loci (*1, 2*). In all animals, many olfactory receptor genes are found in tandem arrays, as a result of their duplication by non-allelic homologous recombination (*6*). In some cases, such clustered receptors share enhancer elements (*1, 2*), but whether this genomic organization has other consequences is unclear. Here, through analysis of mechanisms controlling the expression of a model tandem cluster of three olfactory receptor genes (*Ir75c, Ir75b* and *Ir75a* (organized 5’-3’) (*7*)) in *Drosophila*, we have discovered, serendipitously, novel types of feedback pathway to ensure exclusive receptor expression.

*Ir75c, Ir75b* and *Ir75a* are encoded by a cluster of genes that are thought to have arisen through duplication of an *Ir75a-like* ancestral gene in the last common Drosophilidae ancestor (*7*). These volatile acid-sensing “tuning” receptors are expressed – together with the co-receptor Ir8a (*8*) – in distinct, spatially-stereotyped populations of olfactory sensory neurons (OSNs) in the main olfactory organ, the antenna (Fig. 1A) (*7, 9*). To identify *trans*-acting factors that regulate the expression of these receptor genes, we performed a transgenic RNAi screen of 121 genes encoding transcription factors and chromatin regulators that were previously implicated in peripheral olfactory system development (*10*). We induced RNAi with a constitutive Gal4 driver that is active broadly throughout antennal development (*10*) (Fig. 1B), and examined expression of Ir75c, Ir75b, Ir75a and Ir8a with antibodies, as well as overall antennal morphology.

**Fig. 1.**
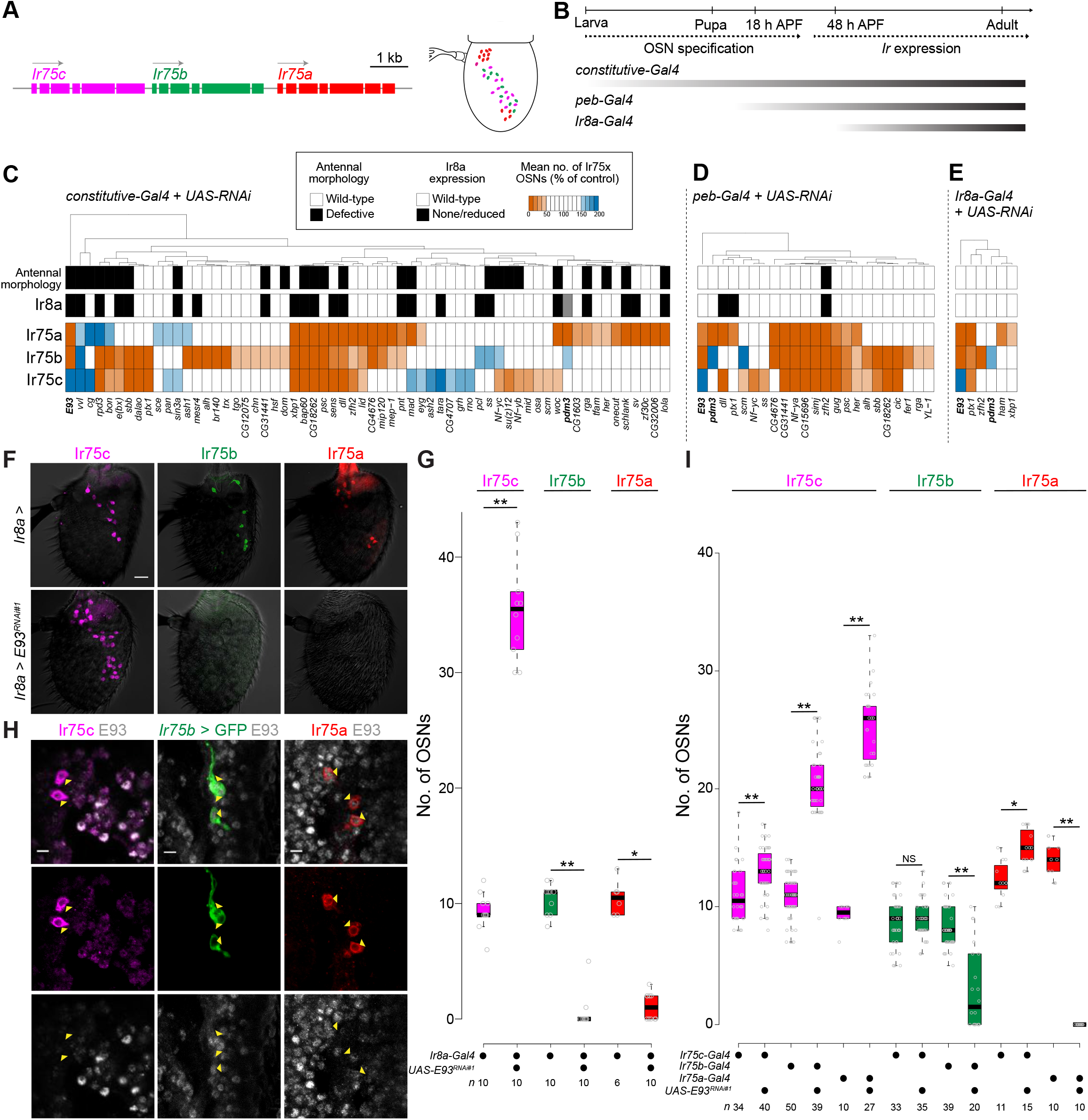
An RNAi screen identifies a dual role for E93 in repression and activation of *Ir* genes. (**A**) Schematics of the tandem array of *Ir75c, Ir75b* and *Ir75a* genes (arrows indicate the direction of transcription) and the distribution of the neurons expressing the corresponding proteins in the antenna. (**B**) Schematic of the main phases of OSN development from the larval antennal disc to the adult antenna; the temporal expression patterns of the three drivers used in this study are shown below (APF, after puparium formation). (**C**) Positive hits of the *constitutive-Gal4* RNAi screen. At least 3 antennae were scored for each line (see Supplementary Table 1 for screen data for (C-D)). For Ir75a/b/c cell numbers, an RNAi line was considered to give a phenotype if there were fewer than half, or more than 1.5×, the numbers of control genotypes (as illustrated by the heat-map). Euclidian distance was used for the hierarchical clustering in (C-E). The grey box indicates the missing data for Ir8a phenotype for *pdm3*. Control genotype: *ey-FLP,Act5c*(*FRT-CD2*)-*Gal4/+*. (**D**) Positive hits of the *peb-Gal4* RNAi screen. Control genotype: *peb-Gal4,UAS-Dcr-2/+*. (**E**) Positive hits of the *Ir8a-Gal4* RNAi screen. Control genotype: *Ir8a-Gal4/+*. (**F**) Representative images of Ir75c, Ir75b and Ir75a immunofluorescence in control (*Ir8a-Gal4/+*) and *E93^RNAi^* (*Ir8a-Gal4/UAS-E93^RNAi#1^*) antennae. Scale bar, 20 μm. The fluorescent channels are overlaid on a brightfield background to show antennal morphology. (**G**) Quantification of neuron numbers in the genotypes shown in (F). Boxplots show the median, first and third quartile of the data, overlaid with individual data points. Pairwise Wilcoxon rank-sum test: ** *P* < 0.001; * *P* < 0.05; NS *P* > 0.05. The sample size is shown beneath the plot. (**H**) Representative images of immunofluorescence on antennal sections for E93 and Ir75c or Ir75a in *w^1118^* flies, or E93 and GFP in *Ir75b-Gal4/+;UAS-myr:GFP/+* animals. Yellow arrowheads indicate the neuron soma. Scale bar, 5 μm. (**I**) Quantification of neuron numbers in control and Ir OSN-targeted *E93^RNAi^* antennae. Genotypes: *UAS-myr:GFP/+;Ir75c-Gal4/+, UAS-myr:GFP/+;Ir75c-Gal4/UAS-E93^RNAi#1^*, *Ir75b-Gal4/+;UAS-myr:GFP/+, Ir75b-Gal4/+;UAS-myr:GFP/UAS-E93^RNAi#1^*, *Ir75a-Gal4/+*;[*UAS-myr:GFP* or +]/+, *Ir75a-Gal4/+*;[*UAS-myr:GFP* or +]/*UAS-E93^RNAi#1^* (GFP signal was not used in any quantifications). Pairwise Wilcoxon rank-sum test: ** *P* < 0.001; * *P* < 0.05; NS *P* > 0.05.

62 genes exhibited RNAi phenotypes (Fig. 1C, Fig. S1A,B, and Supplementary Table 1): most of the phenotypes reflect reductions in the number of OSNs expressing these tuning receptors, but increases in OSN population size were occasionally observed. RNAi phenotypes could encompass the expression all three tuning receptors (and often also Ir8a), but many affected only one OSN type or different combinations of two neuron classes, revealing a complex and unique gene regulatory network for each OSN population.

Because RNAi is induced early in OSN development, many of the tested genes are likely to have upstream roles in antennal development and/or OSN fate specification; indeed many lines produced morphological defects of the antenna (Fig. 1C, Fig. S1B and Supplementary Table 1). To identify candidate molecules that directly regulate receptor expression we performed RNAi on the initial hits using two additional drivers, *pebbled* (*peb*)-*Gal4* (*11*) and *Ir8a-Gal4* (*8*), which are expressed ~16 h after puparium formation (APF) and ~48 h APF, respectively (Fig. 1B). As expected, only a subset of genes showed phenotypes with these drivers (Fig. 1D,E). Notably, co-expression of Ir75c and Ir75b (which were visualized simultaneously) was extremely rare with any driver (Supplementary Table 1), suggesting that redundant mechanisms exist to prevent these receptors from being expressed in the same neuron.

From these screens, we focused on *Ecdysone-induced protein 93F* (*E93*) whose depletion produced a similar phenotype with all drivers: *E93^RNAi^* animals display drastic loss of Ir75a and Ir75b expression, while Ir75c is expressed in many additional neurons (Fig. 1C-G). This phenotype was confirmed in two independent RNAi lines as well as in *E93* mutant animals (Fig. S2A-C). While loss of Ir75a and Ir75b was observed regardless of the driver, the number of Ir75c-expressing cells varied depending upon the timing of knock-down (Fig. S2A).

Previous studies showed that E93 is expressed broadly in the antenna, and is necessary for the expression of several *Odorant receptor* (*Or*) genes (*4*). Co-labeling with E93 and Ir antibodies (or an *Ir* transgenic reporter (*7*)) revealed that E93 is present in Ir75b and Ir75a neurons, but not in Ir75c neurons (Fig. 1H). *Ir8a-Gal4*-driven *E93^RNAi^* (hereafter, *Ir8a>E93^RNAi^*) occurs at the time when tuning receptor transcription begins (*12*), suggesting that E93 has a very proximal (if not direct) role in regulating expression of these receptors. The number and spatial distribution of additional Ir75c-positive cells in *Ir8a>E93^RNAi^* antennae were consistent with Ir75b and Ir75a neurons expressing Ir75c instead of their normal receptors (Fig. 1A,F,G). We tested this possibility by inducing *E93^RNAi^* with drivers for these individual neuron populations (Fig. 1I). *Ir75a>E93^RNAi^* led to loss of Ir75a expression, and increase in Ir75c OSN numbers. *Ir75b>E93^RNAi^* led to reduction (but not complete loss) of Ir75b expression, and increase in Ir75c OSN numbers. *Ir75c>E93^RNAi^* has little phenotypic consequence, consistent with the lack of E93 expression in this neuron population.

Together, these observations suggested an initial simple model: in Ir75a and Ir75b neurons, E93 promotes expression of Ir75a or Ir75b (presumably together with OSN subtype-specific transcription factors) and inhibits expression of Ir75c. In Ir75c neurons, E93 is absent, which permits expression of Ir75c (induced by other transcriptional activators).

In other tissues, E93 regulates local chromatin structure and/or directly controls gene transcription (*13*), raising the question of whether expression of *Ir75c, Ir75b* and *Ir75a* is affected at the transcriptional level by the absence of this protein. The boundaries of transcription units of these receptor genes were previously unclear, confounded by the detection of apparent chimeric transcripts containing exons from adjacent members of this cluster (*7*). To clarify this issue, and determine the effect of loss of E93 on transcription from these loci, we performed bulk RNA-sequencing (RNA-seq) and quantitative reverse transcription-PCR (qRT-PCR) of control and *E93^RNAi^* antennae.

RNA-seq analysis of control tissue provided evidence for the existence of seven distinct transcripts encoded by these three receptor genes (Fig. 2A). Notably, many transcripts initiating from *Ir75c* do not terminate at the 3’ end of this gene but rather run through the *Ir75b* and *Ir75a* exons. These transcripts are very unlikely to encode Ir75b or Ir75a proteins, however, because they lack the first exon (containing the start codon) of these downstream genes. Similarly, a majority of transcripts initiating from *Ir75b* incorporate exons 2-7 of *Ir75a* (Fig. 2A). These extended RNAs appear to reflect a failure in transcription termination at the 3’ end of *Ir75c* and *Ir75b*, consistent with the absence of canonical transcription termination/polyadenylation sequences downstream of their coding regions; such a sequence is only present 3’ of *Ir75a* (Fig. 2A). We confirmed the presence of such transcripts by analysis of Expressed Sequence Tag datasets and Sanger sequencing (data not shown), as well as by RNA fluorescent *in situ* hybridization (FISH; see below and Fig. S3).

**Fig. 2.**
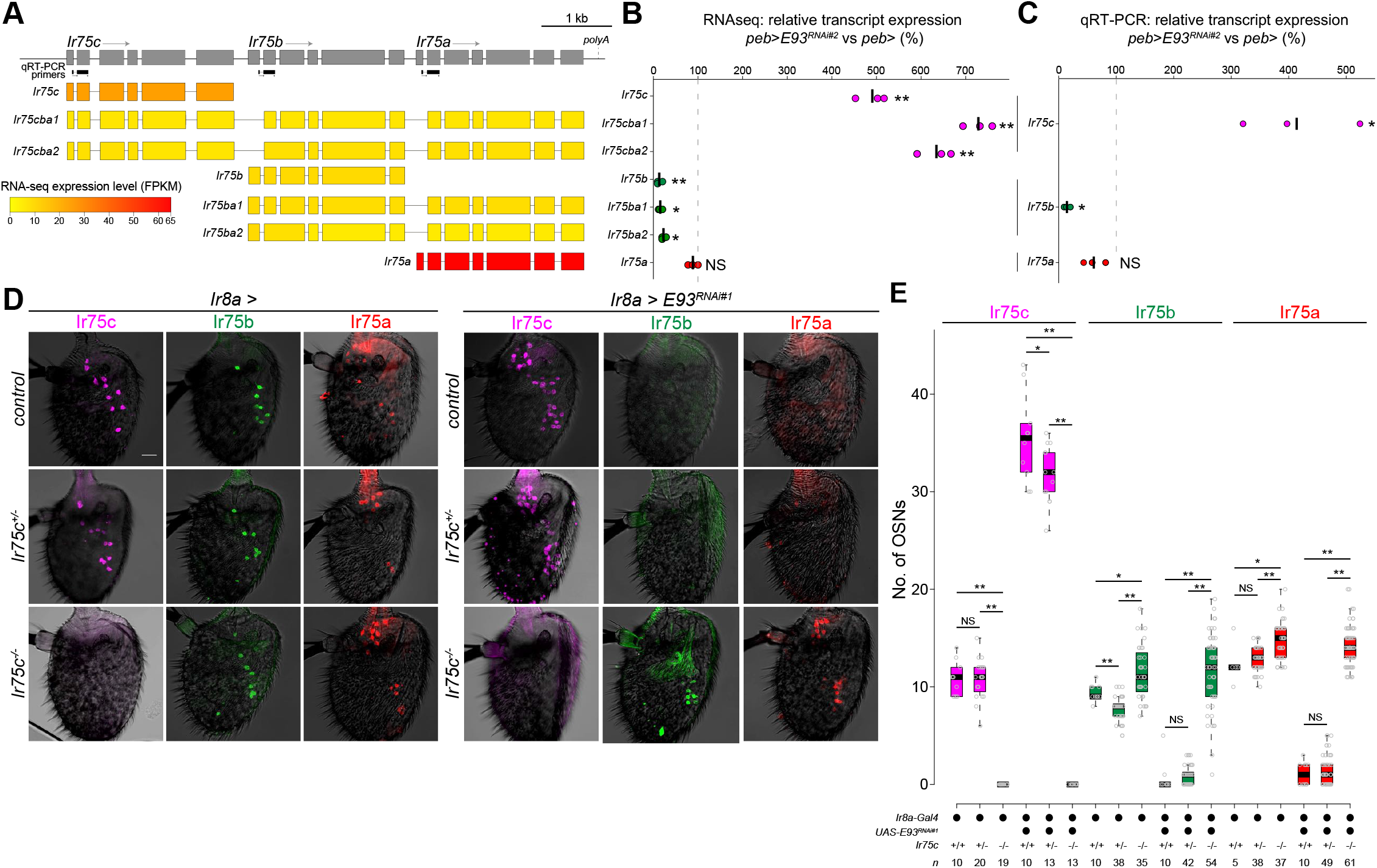
*Ir75c*-dependent repression of Ir75b and Ir75a expression. (**A**) Top: detailed structure of the *Ir75c, Ir75b* and *Ir75a* loci, indicating qRT-PCR primers and the predicted polyadenylation (polyA) site. Bottom: heatmap representation of expression levels of seven isoforms encoded by these genes, determined by RNA-seq of antennae from control flies (*peb-Gal4*,*UAS-Dcr-2/+;VIE-260B/+*). (**B**) Relative expression levels (assessed by RNA-seq) of the seven transcript isoforms in antennae of individual biological replicates (n = 3) of *peb>E93^RNAi#2^* flies (*peb-Gal4,UAS-Dcr-2/+;E93^RNAi#2^/+*) compared to the mean of control samples (genotype as in (A)). The black bar represents the mean. Two sample t-test: ** *P* < 0.001, * *P* < 0.05; NS *P* > 0.05. (**C**) Relative expression (assessed by qRT-PCR) of *Ir75c, Ir75b* and *Ir75a* transcripts in *peb>E93^RNAi#2^* antennae normalized to the mean of the controls (genotypes are as in (A)). The primer pairs used do not distinguish different isoforms for a given gene. Each dot represents the mean value of three technical replicates. Two sample t-test was performed for ddCt values: ** *P* < 0.001, * *P* < 0.05; NS *P* > 0.05 (n = 3 biological replicates). (**D**) Representative images of Ir75c, Ir75b and Ir75a immunofluorescence in antennae of control (left) and *E93^RNAi^* (right) animals in *Ir75c^+/+^, Ir75c^+/−^* or *Ir75c^−/−^* backgrounds. Genotypes: *Ir8a-Gal4/+, Ir8a-Gal4/+;Ir75c^MB08510^/+, Ir8a-Gal4/+; Ir75c^MB08510^/Ir75c^MB08510^, Ir8a-Gal4/UAS-E93^RNAi#1^, Ir8a-Gal4/UAS-E93^RNAi#1^;Ir75c^MB08510^/+, Ir8a-Gal4/UAS-E93^RNAi#1^;Ir75c^MB08510^/Ir75c^MB08510^*. Scale bar, 20 μm. (**E**) Quantification of neuron numbers in the genotypes shown in (D). Pairwise Wilcoxon rank-sum test and P values adjusted for multiple comparisons with the Benjamini and Hochberg method): ** *P* < 0.001, * *P* < 0.05; NS *P* > 0.05.

We next examined changes in transcript levels for the three *Ir* genes in *E93^RNAi^* antennae. Transcripts initiating from *Ir75c* are highly up-regulated in the absence of E93 (Fig. 2B,C), consistent with the ectopic expression of receptor protein (Fig. 1F,G). Coding transcripts for *Ir75b* and *Ir75a* were identified as those containing the first exon of these genes: transcription initiating from *Ir75b* is strongly diminished (Fig. 2B,C), concordant with the loss of Ir75b expression (Fig. 1F,G). By contrast, *Ir75a* transcripts levels were only slightly, but non-significantly, reduced in *E93^RNAi^* antennae (Fig. 2B,C), despite the loss of detectable Ir75a protein (Fig. 1F,G). These observations refined our model as they indicate that E93 inhibits *Ir75c* transcription and promotes *Ir75b* transcription, but has only an indirect role in promoting *Ir75a* expression.

We were unable to identify an E93 binding motif (*13*) within the minimal promoter element of *Ir75b* (*7*). Given the increase in Ir75c expression in Ir75b neurons (Fig. 1I) and the existence of transcriptional read-through across these loci (Fig. 2A), we hypothesized that the decrease in *Ir75b* transcription is an indirect consequence of the ectopic expression of *Ir75c* through a process of transcriptional interference (*14*), *i.e.*, where transcription from one gene (here, *Ir75c*) inhibits, in *cis*, the expression of another (here, *Ir75b*).

To test this possibility, we obtained an *Ir75c* mutant (*Ir75c^MB08510^*) bearing a *Minos* insertion in the 5’ regulatory region (Fig. S3A), which we found leads to essentially complete loss of *Ir75c* RNA (Fig. S3B-D). By contrast, transcripts of *Ir75b* and *Ir75a* were still detected (Fig. S3C). Moreover, analysis of the number of neurons detected by RNA probes for these receptor genes in control and *Ir75c* mutants provided further evidence of read-through of *Ir75c* transcripts through these downstream loci (Fig. S3C-D). We noticed that numbers of Ir75b (and Ir75a) neurons were slightly elevated in *Ir75c* mutant antennae compared to controls (Fig. 2D,E); we return to this phenomenon in the last section of the results. Most strikingly, combination of the *Ir75c* mutant with *E93^RNAi^* was sufficient to restore Ir75b expression (Fig. 2D,E). This result has two implications: first, that E93 is not directly required for transcription of *Ir75b* and second, that expression of *Ir75c* can inhibit transcription of *Ir75b*.

Importantly, loss of *Ir75c* in an *E93^RNAi^* background also led to restoration of Ir75a expression (Fig. 2D,E). As *Ir75a* is still transcribed in *E93^RNAi^* antennae (Fig. 2B,C), this observation indicates that *Ir75c* inhibits *Ir75a* expression post-transcriptionally. Additionally, Ir75b expression is only very slightly increased in *E93^RNAi^* antennae in a heterozygote *Ir75c* mutant (where one allele of *Ir75b* should, in principle, not be subject to transcriptional interference by *Ir75c*) (Fig. 2E), implying that Ir75b must also be repressed post-transcriptionally by Ir75c.

We examined this possibility by misexpressing *Ir75c* from a transgenic cDNA construct (*i.e.*, unlinked to the endogenous *Ir75c* locus) using *Ir8a-Gal4*. We observed a mild but significant reduction in the number of Ir75b-expressing cells and more substantial loss of detectable Ir75a (Fig. 3A,B). Many of the remaining Ir75b and Ir75a neurons had only weak immunofluorescence signals (Fig. 3C). As a control, we misexpressed *Ir75a* cDNA using the same driver, but this had no obvious impact on Ir75c expression, and led to a slight but variable increase in Ir75b (Fig. 3A-C). Together with the analysis of *Ir* transcript levels (Fig. 2B,C), these data are consistent with a model in which *Ir75c* expression inhibits *Ir75b* both at the transcriptional level and post-transcriptional level, while Ir75c inhibits Ir75a expression predominantly post-transcriptionally.

**Fig. 3.**
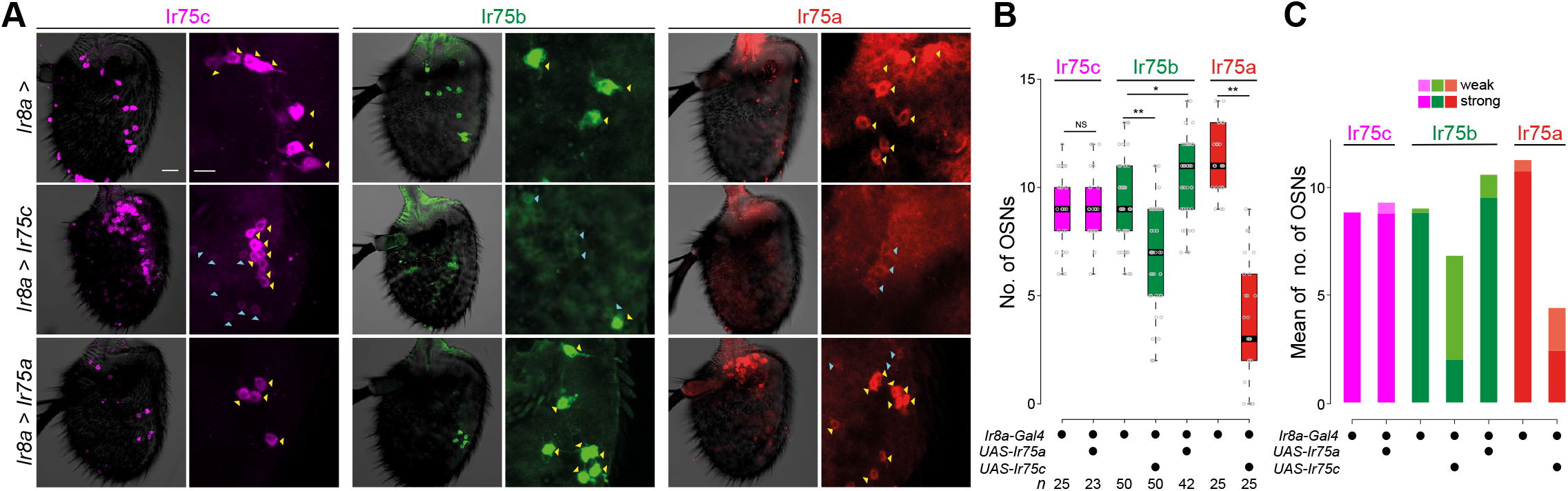
Post-transcriptional inhibition of Ir75a and Ir75b by Ir75c. (**A**) Representative images of Ir75c, Ir75b and Ir75a immunofluorescence in antennae of control (*Ir8a-Gal4/+*), *Ir75c* mis-expression (*Ir8a-Gal4/+;UAS-Ir75c/+*) and *Ir75a* misexpression (*Ir8a-Gal4/+;UAS-Ir75a/+*) animals. Scale bar, 20 μm. A region of interest is shown at higher magnification to the right of each image; scale bar, 10 μm. Yellow and cyan arrowheads indicate neurons with strong and weak receptor expression, respectively. (**B**) Quantification of neuron numbers in the genotypes shown in (A) (Ir75a quantification for *Ir8a>Ir75a* and Ir75c quantification for *Ir8a>Ir75c* antennae are not shown, as neuron density – comprising the entire Ir8a population – is too high for unambiguous scoring). Pairwise Wilcoxon rank-sum test and P values adjusted for multiple comparisons with the Bonferroni method: ** *P* < 0.001, * *P* < 0.05; NS *P* > 0.05. (**C**) Quantification of weak and strong neuron numbers in the genotypes shown in (A).

The failure of *Ir75b* transcripts to terminate efficiently at the 3’ end of this gene (Fig. 2A) raised the possibility that *Ir75b* can also suppress expression of *Ir75a* by transcriptional interference, analogous to the inhibition of *Ir75b* transcription by *Ir75c*. From our RNAi screen, we noted that both *constitutive-Gal4* and *peb-Gal4-driven* RNAi of *pou domain motif 3* (*pdm3*), led to increased numbers of Ir75b-expressing cells and greatly diminished the number of Ir75a-expressing cells (Ir75c expression is unchanged) (Fig. 1C,D). An independent study of Pdm3 indicated that this transcription factor is expressed in Ir75a neurons, but not Ir75b or Ir75c OSNs; here, Pdm3 promotes Ir75a neuron fate and suppresses Ir75b neuron fate, encompassing receptor protein expression and axon targeting properties (*15*). We wondered, however, whether loss of Ir75a in *pdm3^RNAi^* antennae might be an indirect consequence of ectopic Ir75b expression.

We first examined whether loss of *pdm3* affects transcription of these genes. Indeed, qRT-PCR analysis corresponded well with changes in the number of neurons expressing Ir75b and Ir75a proteins: *peb>pdm3^RNAi^* antennae displayed an increase in *Ir75b* transcription and decreased in *Ir75a* transcription (Fig. S4A). Furthermore, we confirmed the increase in number of cells expressing *Ir75b* transcripts by RNA FISH upon loss of Pdm3 (Fig. S4B,C). The number of neurons detected by our probe for *Ir75a* RNA was not different between genotypes, presumably because this reagent detects transcripts initiating from all three genes in the cluster; the decrease in *Ir75a*-expressing cells is therefore counter-balanced by the increase in the size of the Ir75b neuron population (Fig. S4B,C).

To test whether Pdm3’s role in promoting *Ir75a* transcription was direct or indirect, we combined *peb>pdm3^RNAi^* with an *Ir75b* mutant (*Ir75b^DsRed^*) which bears a 1.3 kb insertion in the middle of the gene that abolishes *Ir75b* transcript and protein expression (Fig. S3A,E,F and Fig. 5A,B). Importantly, Ir75a expression is partially restored through loss of one copy of *Ir75b*, and fully restored in homozygous *Ir75b* mutants (Fig. 4A,B). Together, these data support a model in which Pdm3 functions in Ir75a neurons to repress *Ir75b* expression, which would otherwise inhibit transcription of *Ir75a* in *cis*. We tested whether Ir75b might also impact Ir75a expression post-transcriptionally, but found *Ir8a*-*Gal4*-driven heterologous expression of a *UAS-Ir75b* transgene yielded barely detectable ectopic Ir75b (data not shown), precluding conclusive insights.

**Fig. 4.**
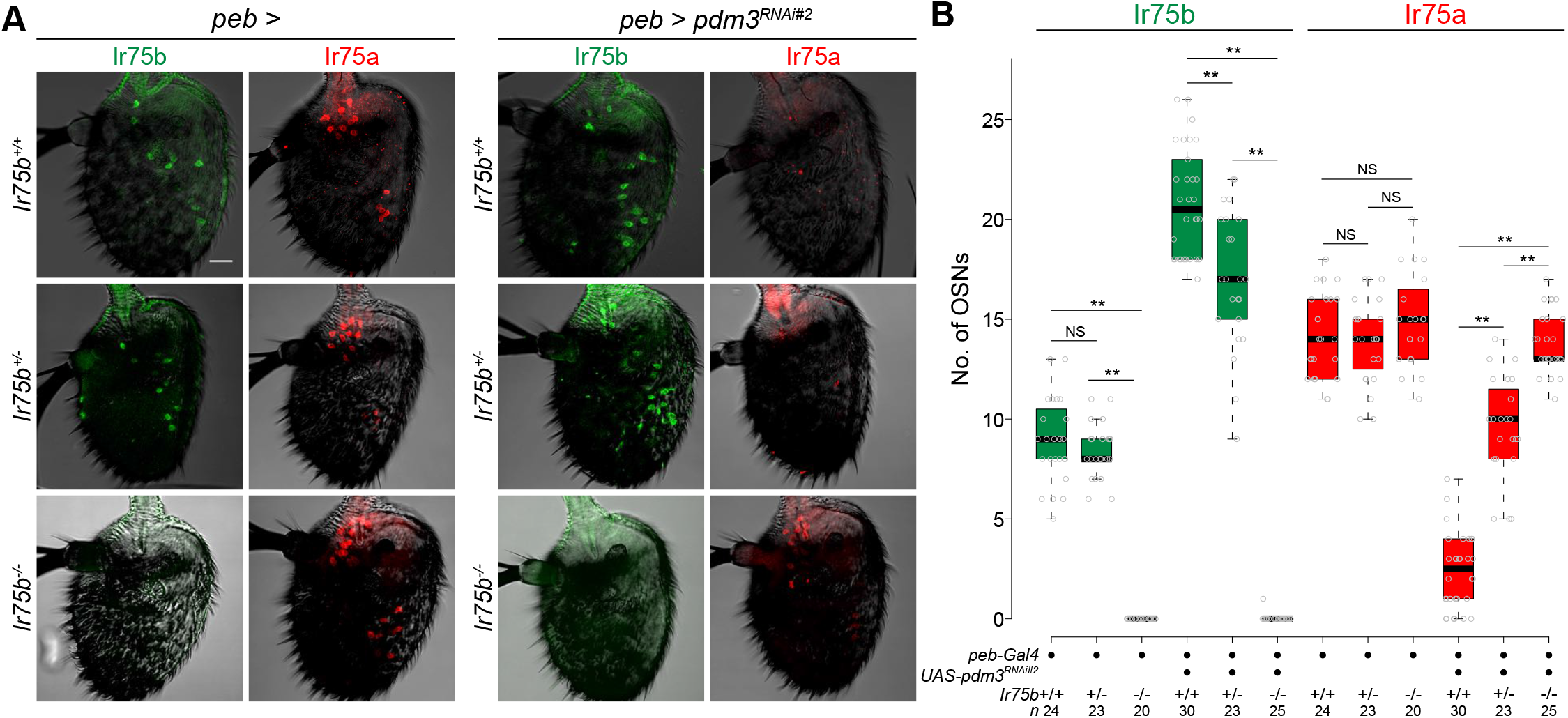
*Ir75b*-dependent repression of *Ir75a*. (**A**) Representative images of Ir75b and Ir75a immunofluorescence in control and *pdm3^RNAi^* antennae in *Ir75b^+/+^, Ir75b^+/−^* and *Ir75b^-/-^* backgrounds. Genotypes (top to bottom): *peb-Gal4,UAS-Dcr-2/+, peb-Gal4,UAS-Dcr-2/+;;Ir75b^DsRed^/+, peb-Gal4, UAS-Dcr-2/+;Ir75b^DsRed^/ Ir75b^DsRed^* (left), *peb-Gal4, UAS-Dcr-2/+;UAS-pdm3^RNAi^#^2^/+, peb-Gal4, UAS-Dcr-2/+;UAS-pdm3^RNAi#2^/+;Ir75b^DsRed^/+, peb-Gal4,UAS-Dcr-2/+;UAS-pdm3^RNAi#2^/+;Ir75b^DsRed^/Ir75b^DsRed^* (right). Scale bar, 20 μm. (**B**) Quantification of neuron numbers in the genotypes shown in (A). Pairwise Wilcoxon rank-sum test and P values adjusted for multiple comparisons with the Bonferroni method: ** *P* < 0.001, * *P* < 0.05; NS *P* > 0.05.

**Fig. 5.**
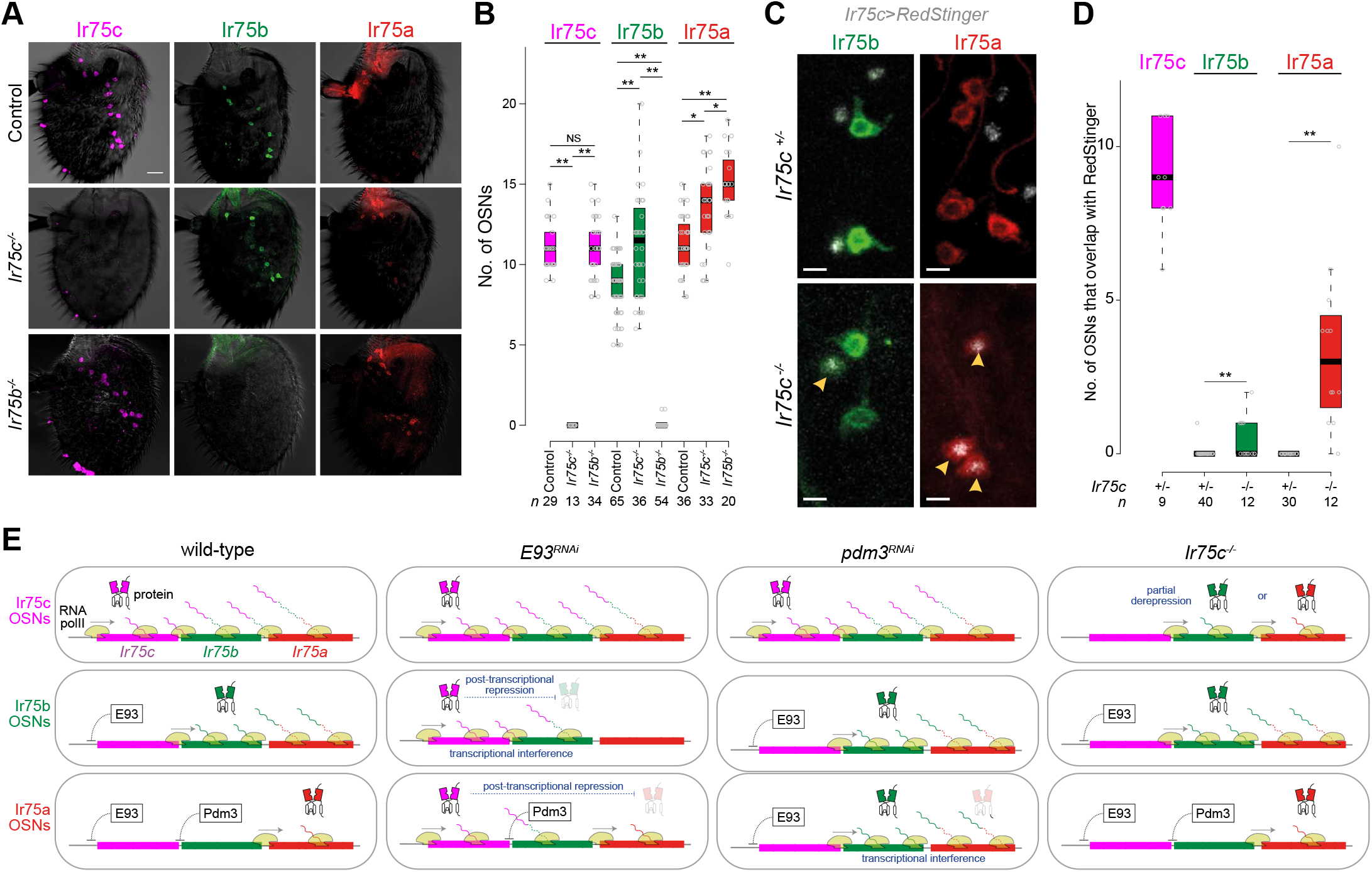
Ectopic expression of Ir75b and Ir75a in *Ir75c* mutant neurons. (**A**) Representative images of Ir75c, Ir75b and Ir75a immunofluorescence in control (*w^1118^*), *Ir75c^-/-^* and *Ir75b^-/-^* antennae. Scale bar, 20 μm. (**B**) Quantification of neuron numbers in the genotypes shown in (A). Pairwise Wilcoxon rank-sum test and P values adjusted for multiple comparisons with the Benjamini and Hochberg method: ** *P* < 0.001, * *P* < 0.05; NS *P* > 0.05. (**C**) Representative images of Ir75b and Ir75a immunofluorescence for a subset of Ir75c neurons (labelled with RedStinger (grey)) in *Ir75c^+/-^* and *Ir75c^-/-^* flies are shown. Yellow arrowheads mark *Ir75c* mutant neurons that ectopically express Ir75a or Ir75b. Genotypes: *UAS-RedStinger/+;Ir75c-Gal4,Ir75c^MB08510^/+, UAS-RedStinger/+;Ir75c-Gal4,Ir75c^MB08510^/Ir75c-Gal4,Ir75c^MB08510^*. Scale bar, 5 μm. (**D**) Quantification of Ir75c neurons that ectopically express Ir75b or Ir75a in *Ir75c^+/-^* and *Ir75c^-/-^* flies (genotypes as in (C)). Pairwise Wilcoxon rank-sum test and P values adjusted for multiple comparisons with the Bonferroni method: ** *P* < 0.001, * *P* < 0.05; NS *P* > 0.05. In the control Ir75c immunofluorescence samples, all neurons (81/81 (100%)) express both Ir75c and *Ir75c>RedStinger*. (**E**) Model of the regulatory interactions between *Ir75c, Ir75b* and *Ir75a* gene products in wild-type, *E93^RNAi^, pdm3^RNAi^* and *Ir75c^-/-^* genetic backgrounds for Ir75c, Ir75b and Ir75a neurons. Individual transcripts and long transcripts lacking the first exons of the downstream genes are shown as unbroken and broken lines, respectively. It is unknown whether E93 and Pdm3 inhibit transcription of *Ir75c* and *Ir75b*, respectively, directly or indirectly (indicated by dashed lines).

Finally, we asked whether Ir75c and Ir75b exert a repressive effect on other genes in the cluster in their endogenous neurons, and not only when expressed ectopically in other neurons (as in *E93^RNAi^* or *pdm3^RNAi^* antennae or through transgenic misexpression of Ir75c). In *Ir75c* mutant antennae, we observed a small and variable increase in the number of neurons expressing Ir75b and Ir75a, compared to controls (Fig. 5A,B; see also Fig. 2E). Similarly, in *Ir75b* mutants, we observed an increase in numbers of neurons expressing Ir75a, but no effect on those expressing Ir75c, consistent with a contribution of Ir75b in suppressing expression of Ir75a in its own neurons (Fig. 5A,B). This latter effect was sensitive to the genetic background, however, as we did not observe such an increase in all genetic backgrounds (e.g., Fig. 4B).

We therefore focused on the *Ir75c* mutant, and examined whether the additional expression of Ir75b and Ir75a is in Ir75c neurons, by labeling these cells transgenically with *Ir75c>RedStinger*. In control tissue we detected, as expected, no expression of Ir75a or Ir75b in Ir75c neurons (with the exception of one cell) (Fig. 5C,D). By contrast, in *Ir75c* mutants, several Ir75c neurons now express these Irs (Fig. 5C,D), indicating that Ir75c represses these receptors in its own neural population. It is not surprising that the ectopic expression of Ir75b and Ir75a is limited to only a small subset of *Ir75c* mutant neurons as these cells presumably only partially resemble the gene regulatory environment of endogenous Ir75b or Ir75a neurons, as illustrated by our original RNAi screens (Fig. 1C-E).

This work has identified many new candidate activators and repressors of *Ir* gene transcription. Importantly, analysis of two of these, E93 and Pdm3, has revealed novel mechanisms by which clustered receptor genes can repress each other to contribute to their unique expression patterns (Fig. 5E). These discoveries provide, to our knowledge, the first evidence for inter-olfactory receptor locus regulation in invertebrates.

The *cis*-mediated transcriptional repression of *Ir75b* by *Ir75c*, and of *Ir75a* by *Ir75b*, is most simply explained by transcriptional interference (*14*). Beyond the existence of transcripts spanning exons from two or more genes within this array, our previous mapping of RNA polymerase II occupancy in these neurons by Targeted DamID (*15*) provides additional evidence for the failure in efficient termination of the transcription machinery at the 3’ end of *Ir75c* and *Ir75b*. Several *Gustatory receptor* (*Gr*) genes are arranged in comparably tight clusters, with transcription termination signals only apparent after the last gene in the array (*16*). Analysis of transcripts encoded by the clustered sugar-sensing *Gr* genes reveals that they encompass exons of several individual loci (*16, 17*). While these observations were previously interpreted as evidence for polycistronic transcription of these genes (*16, 17*), it is possible that such transcripts reflect a mechanism by which *Gr* genes impact each other’s expression. The existence of transcriptional interference in recently-duplicated chemosensory genes may also provide a substrate for natural selection to favor rapid acquisition of distinct expression patterns – or otherwise pseudogenization of one duplicate – as suggested by genome-wide expression analysis of nested (non-chemosensory) genes in drosophilids (*18*). In this context, E93 and Pdm3 fulfill important inhibitory roles in these neurons’ gene regulatory networks: by preventing transcription of upstream genes in the cluster, they permit expression of downstream genes (Fig. 5E).

The mechanism of post-transcriptional (*trans*) suppression of Ir75b and Ir75a by Ir75c remains unclear. As *Ir75a* transcription is not substantially diminished by *Ir75c* misexpression, it seems unlikely that it operates via the same feedback mechanism acting in mammals, which ultimately ensures the silenced transcriptional state of non-chosen receptor genes (*2*). One hypothesis is that Ir protein subunits compete with each other for the common Ir8a subunit, which is essential for their stabilization in OSNs (*8, 19*). However, combined misexpression of Ir8a with Ir75c did not counter the loss of Ir75b and Ir75a expression (data not shown), suggesting that this co-receptor is not a limiting factor. It is also formally possible that the *Ir75c* transcripts impede translation of other *Ir* transcripts; this is not likely to occur via antisense inhibition, as we have not detected evidence for such transcripts in our bulk RNA-seq data. Precise details of the repressive mechanism(s) will require future development of methods to assay and quantify the processes leading to expression of receptor transcripts and proteins within specific OSN populations.

Given the widespread occurrence of chemosensory receptor families in arrays in the genomes of all animals (*6*), our discoveries are likely to have broad implications for the evolution of their distinct expression patterns. Moreover, the dominance of expression of 5’ located *Ir* genes over 3’ located genes is reminiscent of the “posterior dominance” and “posterior prevalence” of mRNA levels and protein activity observed for clustered *Hox* transcription factor genes (*20, 21*). Study of the interactions between grouped olfactory receptor genes may reveal general insights into how other types of proteins encoded by tandemly-duplicated genes evolve distinct expression patterns.

## Acknowledgements

We thank Eric Baehrecke and Chris Doe for reagents, Christopher Dumayne for help with qRT-PCR experiments, and Maria Cristina Gambetta and members of the Benton laboratory for comments on the manuscript. L.L.P.-G. was supported by a FEBS Long-Term Fellowship. T.O.A. was supported by a Human Frontier Science Program Long-Term Fellowship (LT000461/2015-L). Research in R.B.’s laboratory is supported by the University of Lausanne, ERC Consolidator and Advanced Grants (615094 and 833548, respectively) and the Swiss National Science Foundation.

## Author contributions

All authors contributed to experimental design, analysis and interpretation of results. K.M. performed all experiments, except for those in Fig. S3C-F and Fig. S4B,C, as well as RNA isolation for RNA-seq and qRT-PCR experiments, which were performed by S.C. P.C.C. provided the candidate gene list for the RNAi screen and performed initial analysis of RNA-seq data. L.L.P.-G. supervised K.M. during initial stages of the project. T.O.A. generated the *Ir75b* mutant. S.P. performed RNA-seq data analysis. K.M. and R.B. wrote the paper with input from all other authors.

## Methods

### *Drosophila* strains

Flies were maintained at 25°C in 12 h light:12 h dark conditions, except where noted. *D. melanogaster* strains were maintained on a standard wheat flour-yeast-fruit juice medium. Published mutant and transgenic *D. melanogaster* are described Supplementary Table 2. For histological experiments, flies were 3-8 days old. To increase the efficiency of knockdown, flies were moved to 27°C after 2-3 days. As *peb>E93^RNAi#1^* flies were not viable at 27°C, this genotype was raised at 19°C.

### *Ir75b* mutant generation

The sgRNA expression vector was generated by cloning an annealed oligonucleotide pair (GTCGCCCTCAGTGGCTGACGCGCC and AAACGGCGCGTCAGCCACTGAGGG) into *BbsI*-digested *pCFD3-dU6-3gRNA* (Addgene no. 49410), as described (*22*). The donor vector was constructed by amplifying *Ir75b*-specific homology arms (1.1-1.4 kb) from genomic DNA of *D. melanogaster* Research Resource Identifier Database:Bloomington *Drosophila* Stock Center [RRID:BDSC]_58492 (*23*) (homology arm 1 (5’-3’): GATCCACCTGCGATCTCGCCACCTCGAAGCAGAGCCGGT and GATCCACCTGCGATCCTACCGGCCTGGTAACCCAATCTA; homology arm 2: GATCGCTCTTCGTATGCGTCAGCCACTAAGGGATATGTT and GATCGCTCTTCGGACCACGGCCACCGTGTGATTATGCTG), and inserting the resultant products into *pHD-DsRed-attP* (Addgene no. 51019) (*24*) via *AarI/SapI* restriction cloning.

Transgenesis of *D. melanogaster* was performed in-house following standard protocols (http://gompel.org/methods). For CRISPR/Cas9-mediated homologous recombination, we injected a mix of the sgRNA expression (final concentration 150 ng μl^−1^) and donor vector (final concentration 400 ng μl^−1^) into *D. melanogaster* RRID:BDSC_58492. *Ir75b* mutant flies were selected based on DsRed expression and integration was confirmed by PCR. The *Ir75b^DsRed^* flies were outcrossed to *w^1118^* flies for three generations prior to generating homozygous stocks.

### Histology and image analysis

Immunofluorescence and RNA FISH on whole mount antennae or antennal cryosections were performed essentially as described (*25*). Primary and secondary antibodies used are listed in Supplementary Table 3. Guinea pig anti-Ir75c antibodies were raised against the same peptide epitope (KEYLSELHLRPRLQHRMD) as used to generate the rabbit anti-Ir75c antibodies (*7*) and affinity-purified by Proteintech Group, Inc. RNA FISH probes for *Ir75a* (referred to here as the “*Ir75a* exon probe”), *Ir75b (“Ir75b* exon probe”) and *Ir75c* were as described (*7*). Imaging was performed on a Zeiss confocal microscope LSM710 or, for RNA FISH samples, a Zeiss confocal microscope 880 Airyscan, using 40× or 63× oil immersion objectives. All images were processed in Fiji and the cell counter plugin was used to quantify neuron numbers.

### RNA-sequencing and analysis

Antennal RNA was extracted from three biological replicates of control (*peb-Gal4,UAS-Dcr-2/+;VIE-260B/+* (*VIE-260B* is an empty landing site used for RNAi transgene insertion)) and *E93^RNAi^* (*peb-Gal4,UAS-Dcr-2/+;UAS-E93^RNAi#2/+^*) animals. For each pair of biological replicates, ~350 females were grown under identical conditions and antennae harvested and RNA extracted from 2-5 day-old flies in parallel, as described (*10*). RNA quality was assessed on a Fragment Analyzer (Advanced Analytical Technologies, Inc.); all RNAs had an RQN of 9-10. mRNA isolation, RNA-seq library preparation and sequencing were performed as described (*26*).

Reads were aligned against *Drosophila melanogaster.BDGP6.92* genome using STAR v2.5.3a (*27*). The reads mapped to the region 3L:17,815,000-17,823,000 containing the *Ir75c*/*Ir75b*/*Ir75a* genes were extracted using samtools v1.8 (*28*) and transcripts were assembled using StringTie v1.3.3b (*29*) with parameters for stranded library and a minimum transcript length of 1000 bases. Transcript abundance was quantified with StringTie using the *Drosophila melanogaster*.BDGP6.92 reference annotation complemented with the four novel isoforms (options: -G, -m 1000, -e, -B).

### Quantitative RT-PCR

For analysis of *E93^RNAi^*, the same RNA samples were used as for the RNA-seq experiment. For additional experiments, RNA was isolated from 2-5 day-old flies (~50 antennae) with QIAGEN RNeasy Mini kit (Cat. no. 74104) by following the manufacturer’s guidelines. cDNA was prepared using Invitrogen SuperScript II Reverse Transcriptase (Cat. no. 18064022).

qRT-PCR was performed on an ABI 7500 Real-Time PCR System (Applied Biosystems, Foster city, CA) using iQ SYBR® Green Supermix from Bio-Rad. Primer pairs (5’-3’) were as follows: *Ir75c* (TGCTGGTCCATAAAGGAGGC / GGCTTTGTCGCAACCCAAAT), *Ir75b* (ATTTTGCACTGCTCACTTTCGC / AACTGTACCACTTTTGTCGCA), *Ir75a* (GCTGGCAGAGTGATGAAAGTCT / TCCTGAGTCTGATCGCACTTT) and control (*tubulin: αTub84B isoform*) (TGTCGCGTGTGAAACACTTC / AGCAGGCGTTTCCAATCTG). qRT-PCR was performed with three technical replicate and three biological replicate for each primer pair. The mean of the three technical replicates was used for each biological replicate. ddCt values were calculated with normalization to *tubulin* values and were used for the statistical analysis.

### Statistics and reproducibility

Statistical analyses and plotting were made in RStudio v3.5.2 (R Foundation for Statistical Computing, Vienna, Austria, 2005; R-project-org). For multiple comparisons, a two-tailed Wilcoxon-rank sum test was performed followed by the Benjamini and Hochberg method or the Bonferroni method for P-value adjustment. For Fig. S4A, P value was calculated by one-way ANOVA test of ddCt values followed by Tukey’s post hoc test for multiple comparison. For other qRT-PCR experiments and relative expression of RNA-seq experiment, P values were calculated by two-sample t-test.

### Data availability

All relevant data supporting the findings of this study are available from the corresponding author on request. RNA-seq data will be made available in GEO (Accession GSE150296).

---

**Fig. S1.**
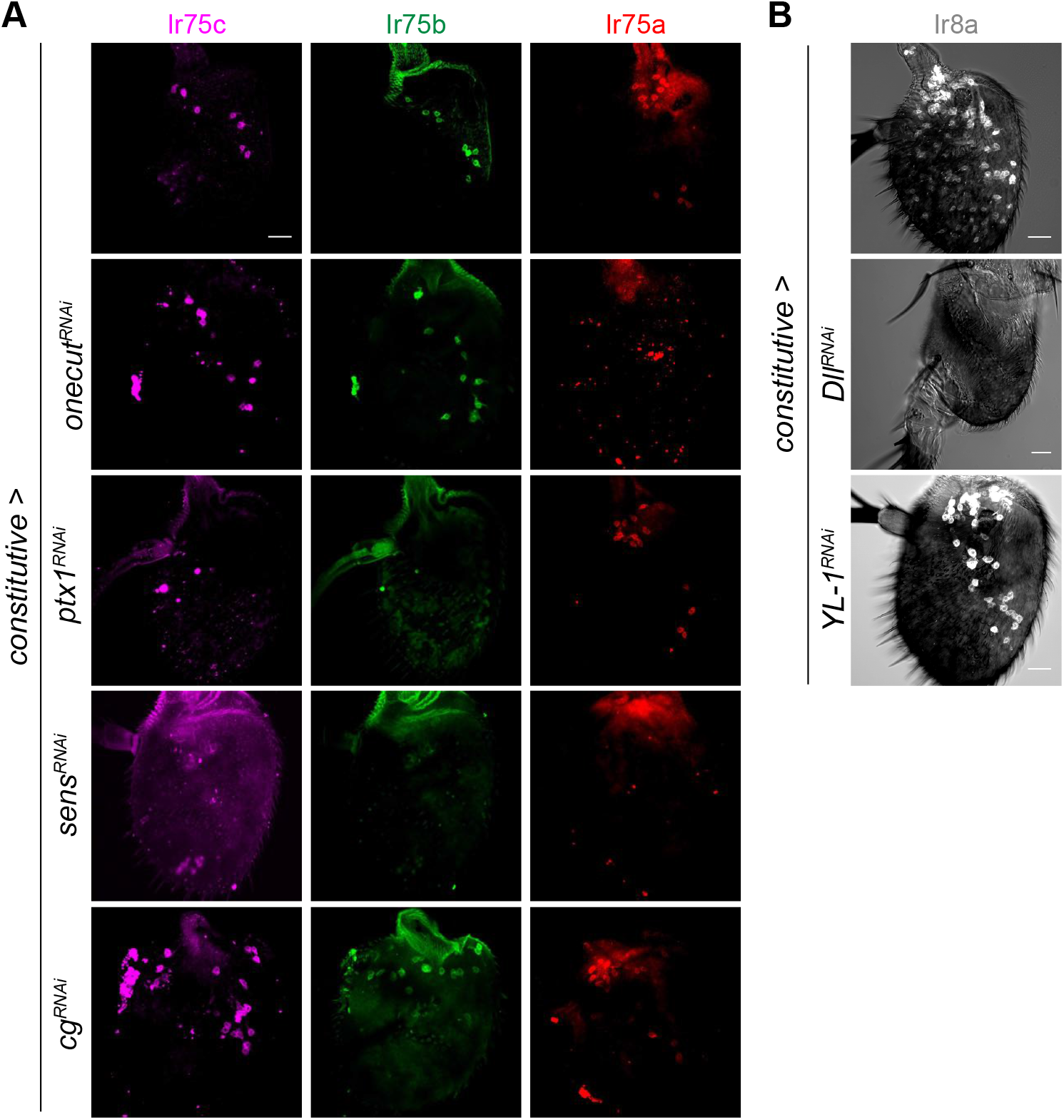
Representative images of a subset of phenotypes from the RNAi screens. (**A**) Representative images of Ir75c, Ir75b and Ir75a immunofluorescence in control and RNAi antennae of the indicated genotypes. Scale bar, 20 μm. (**B**) Representative images of antenna morphology and Ir8a immunofluorescence in control and RNAi antennae of the indicated genotypes. Scale bars, 20 μm.

**Fig. S2.**
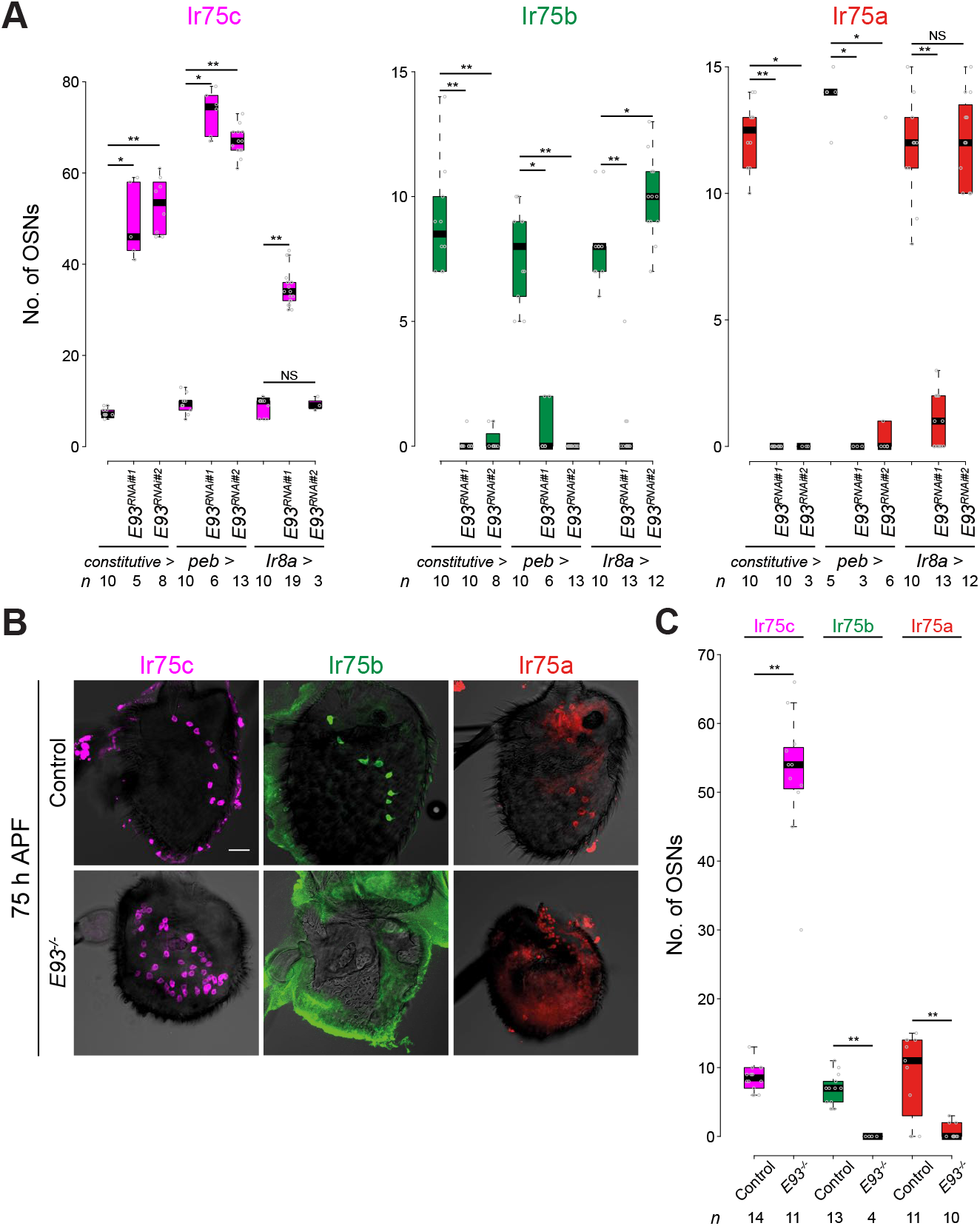
Characterization of *E93* loss-of-function phenotypes. (**A**) Quantification of Ir75c, Ir75b and Ir75a neuron number in control, *E93^RNAi#1^* and *E93^RNAi#2^* flies with *constitutive-Gal4, peb-Gal4* and *Ir8a-Gal4*. Pairwise Wilcoxon rank-sum test and P values adjusted for multiple comparisons with the Benjamini and Hochberg: ** *P* < 0.001, * *P* < 0.05; NS *P* > 0.05. The two RNAi lines were consistent across all drivers and phenotypes, except for *Ir8a-*Gal4-driven *UAS-E93^RNAi#2^*, which did not lead to a phenotype, possibly because of lower RNAi induction. Some of these data incorporated samples from Fig. 1C-G. (**B**) Representative images of Ir75c, Ir75b and Ir75a immunofluorescence of pupal antennae (75 h APF) from control and *E93^-/-^* (*E93^4^/E93^4^*) animals (the mutant animals die just before adult emergence (*30*)). Scale bar, 20 μm. (**C**) Quantification of neuron numbers in the genotypes shown in (B). Pairwise Wilcoxon rank-sum test; ** *P* < 0.001; * *P* < 0.05; NS *P* > 0.05.

**Fig. S3.**
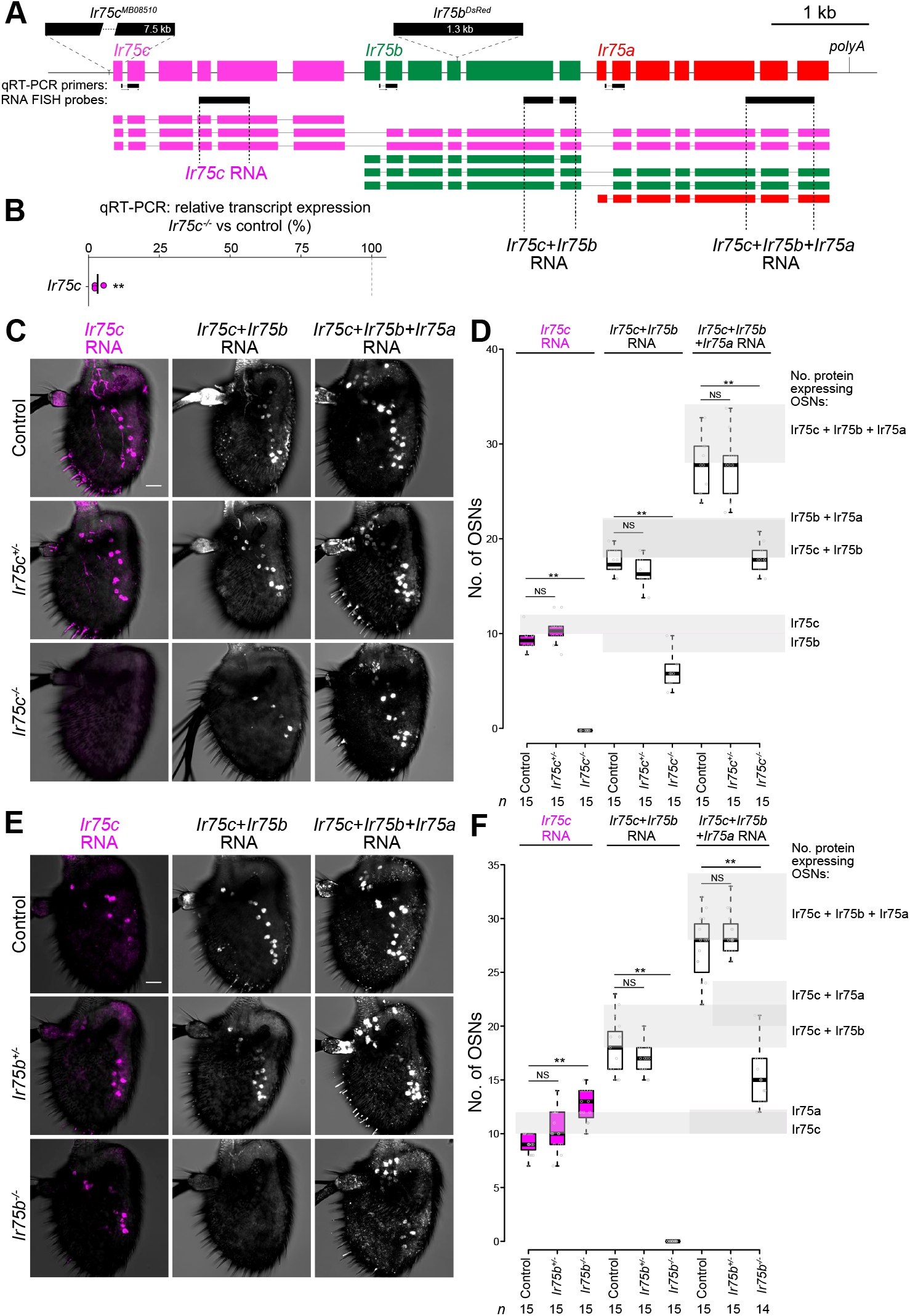
Characterization of *Ir75c* and *Ir75b* mutants. (**A**) Top: structure of the *Ir75c, Ir75b* and *Ir75a* genes indicating the *Ir75c* and *IR75b* mutations and sequences used to generate RNA FISH probes (black bars). Bottom: transcript isoforms (as in Fig. 2A), colored according to the site of initiation (and receptor protein encoded). Gene-specific probes for *Ir75b* and *Ir75a* could not be generated – probes corresponding to the unique short first exons did not give any signal (data not shown) – so the probes covering exons of these genes (referred to as the “*Ir75b* exon probe” and “*Ir75a* exon probe”) also recognize transcripts initiating from upstream genes in the cluster. (**B**) Relative expression (assessed by qRT-PCR) of *Ir75c* transcripts in *Ir75c^-/-^* antennae normalized to the mean of the control (*w^1118^*). Each dot represents the mean value of 3 technical replicates. Two sample t-test of ddCt values: ** *P* < 0.001, * *P* < 0.05; NS *P* > 0.05 (n = 3 biological replicates). (**C**) Representative images of RNA FISH for *Ir75c* (*Ir75c* probe), *Ir75c+Ir75b* (*Ir75b* exon probe) and *Ir75c+Ir75b+Ir75a* (*Ir75a* exon probe) transcripts in control, *Ir75c^+/-^* and *Ir75c^-/-^* antennae. (**D**) Quantification of neuron numbers in the genotypes shown in (C). Grey background shading illustrates the range (between first and third quartiles) of the respective numbers of neurons expressing the indicated receptor proteins (estimated from data in Fig. 5B); these numbers shows good correspondence with the totals observed by RNA FISH in various genotypes (taking into consideration the different sensitivity of RNA FISH and immunofluorescence): in control and *Ir75c^+/-^* antennae, the *Ir75b* exon probe labels Ir75b and Ir75c neurons, and the *Ir75a* exon probe labels Ir75c, Ir75b and Ir75a neurons. By contrast, in *Ir75c^-/-^* antennae, no long (or short) transcripts initiate from *Ir75c*, therefore the *Ir75b* exon probe recognizes only Ir75b neurons, and the *Ir75a* exon probe recognizes only Ir75b+Ir75a neurons. Pairwise Wilcoxon rank-sum test and P values adjusted for multiple comparisons with the Bonferroni method: ** *P* < 0.001, * *P* < 0.05; NS *P* > 0.05. (**E**) Representative images of RNA FISH for *Ir75c* (*Ir75c* probe), *Ir75c+Ir75b (Ir75b* exon probe) and *Ir75c+Ir75b+Ir75a* (*Ir75a* exon probe) transcripts in control, *Ir75b^+/-^* and *Ir75b^-/-^* antennae. (**F**) Quantification of neuron numbers in the genotypes shown in (E). The total number of *Ir75c*-expressing neurons is similar to Ir75c neurons. All labeling by the *Ir75b* exon probe is lost in *Ir75b^-/-^*, as expected (the probe sequence is downstream of the mutational insertion (see (C)). Moreover, in *Ir75b^-/-^* antennae the total number of neurons that contain *Ir75a* RNA decrease to the total number of Ir75a OSNs as in Fig. 5B (slightly higher than control) from the total number of Ir75a+Ir75b+Ir75c OSNs (control and *Ir75b^+/-^* antennae). Pairwise Wilcoxon rank-sum test and P values adjusted for multiple comparisons with the Bonferroni method: ** *P* < 0.001, * *P* < 0.05; NS *P* > 0.05.

**Fig. S4.**
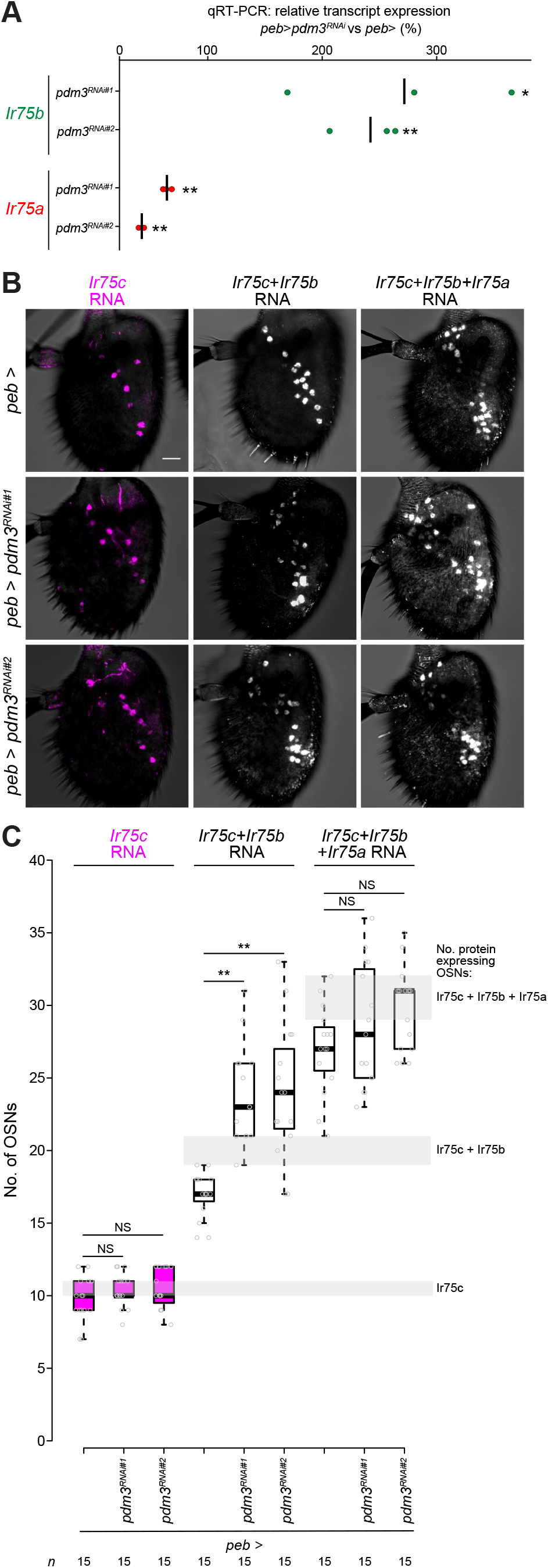
Characterization of *Ir* expression in *pdm3^RNAi^* antennae. (**A**) Relative expression (assessed by qRT-PCR) of *Ir75b* and *Ir75a* transcripts in *pdm3^RNAi^* antennae normalized to control (primers pairs as indicated in Fig. 2A). Genotypes: *peb-Gal4, UAS-Dcr-2/+, peb-Gal4, UAS-Dcr-2/+;UAS-pdm3^RNAi#1^/+, peb-Gal4,UAS-Dcr-2/+;UAS-pdm3^RNAi#2^/+*. Each dot represents the mean value of 3 technical replicates. P values determined by one-way ANOVA test of ddCt values followed by Tukey’s post hoc test for multiple comparison: ** *P* < 0.001, * *P* < 0.05; NS *P* > 0.05 (n = 3 biological replicates). (**B**) Representative images of RNA FISH for *Ir75c* (*Ir75c* probe), *Ir75c+Ir75b* (*Ir75b* exon probe) and *Ir75c+Ir75b+Ir75a* (*Ir75a* exon probe) transcripts for the genotypes in (A). (**C**) Quantification of neuron numbers in the genotypes shown in (A-B). Pairwise Wilcoxon rank-sum test and P values adjusted for multiple comparisons with the Bonferroni method: ** *P* < 0.001, * *P* < 0.05; NS *P* > 0.05. Grey background shows the range (between first and third quartiles) of the respective receptor protein-expressing neuron numbers (estimated from data in Fig. 5B).

**Supplementary Table 1. RNAi screen results.**

Genes, RNAi lines and phenotypes for the screens with the three *Gal4* drivers. If available, more than one RNAi line was used per gene. In some cases, different RNAi lines for a given gene were used with different drivers, due to lethality of certain driver/RNAi line combinations. IF, immunofluorescence. (*provided as a separate Excel file*)

**Supplementary Table 2.**
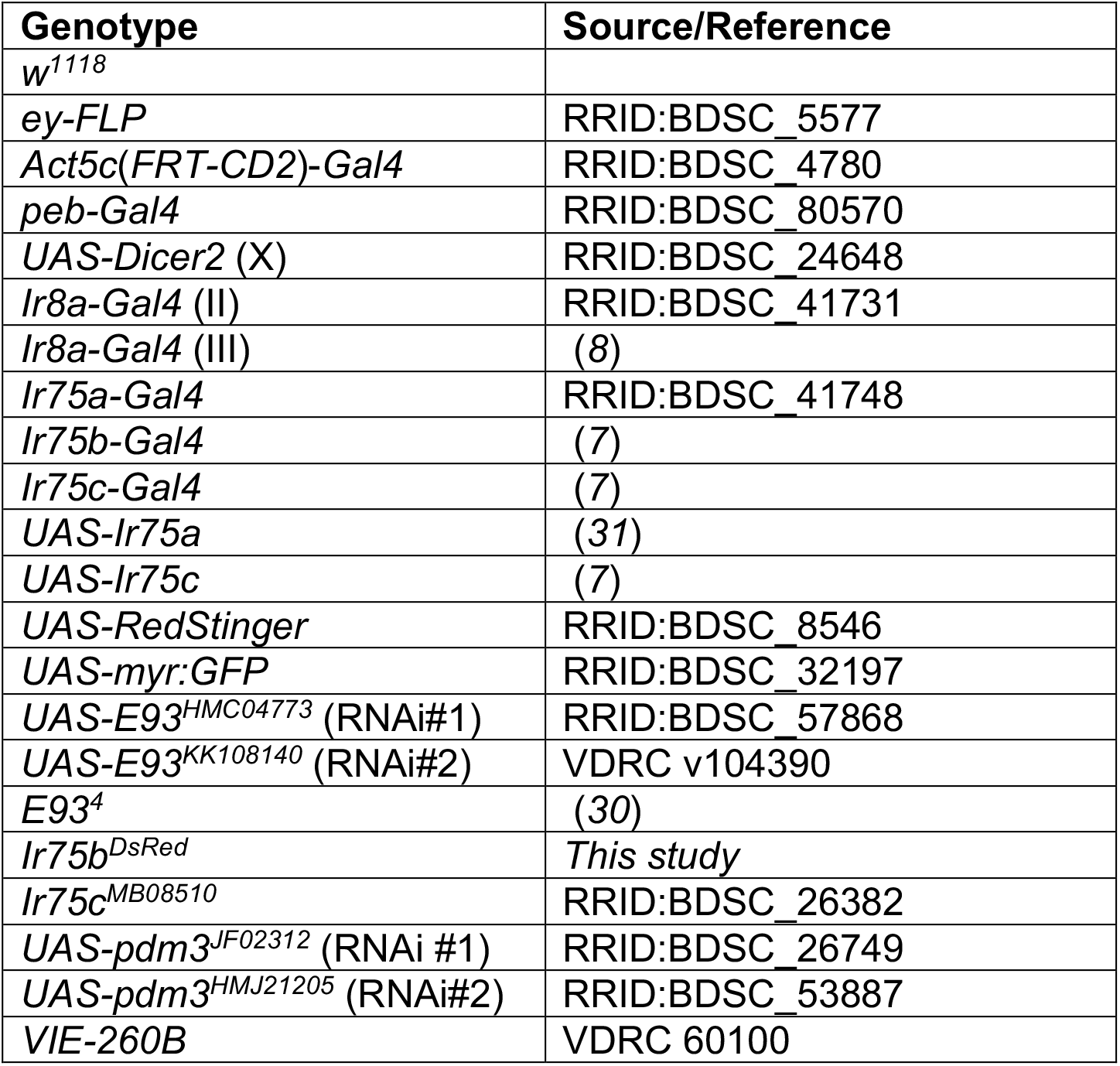
*Drosophila melanogaster* stocks

| Genotype | Source/Reference |
| --- | --- |
| *w<sup>1118</sup>* |  |
| *ey-FLP* | RRID:BDSC_5577 |
| *Act5c(FRT-CD2)-Gal4* | RRID:BDSC_4780 |
| *peb-Gal4* | RRID:BDSC_80570 |
| *UAS-Dicer2* (X) | RRID:BDSC_24648 |
| *Ir8a-Gal4* (II) | RRID:BDSC_41731 |
| *Ir8a-Gal4* (III) | (8) |
| *Ir75a-Gal4* | RRID:BDSC_41748 |
| *Ir75b-Gal4* | (7) |
| *Ir75c-Gal4* | (7) |
| *UAS-Ir75a* | (31) |
| *UAS-Ir75c* | (7) |
| *UAS-RedStinger* | RRID:BDSC_8546 |
| *UAS-myr:GFP* | RRID:BDSC_32197 |
| *UAS-E93<sup>HMC04773</sup>* (RNAi#1) | RRID:BDSC_57868 |
| *UAS-E93<sup>KK108140</sup>* (RNAi#2) | VDRC v104390 |
| *E93<sup>4</sup>* | (30) |
| *Ir75b<sup>DsRed</sup>* | *This study* |
| *Ir75c<sup>MB08510</sup>* | RRID:BDSC_26382 |
| *UAS-pdm3<sup>JF02312</sup>* (RNAi #1) | RRID:BDSC_26749 |
| *UAS-pdm3<sup>HMJ21205</sup>* (RNAi#2) | RRID:BDSC_53887 |
| *VIE-260B* | VDRC 60100 |

**Supplementary Table 3.** Antibodies

| Antibody | Dilution | Source/Reference |
| --- | --- | --- |
| Rabbit anti-Ir75a | 1:100 | RRID:AB_2631091 |
| Guinea pig anti-Ir75b | 1:500 | RRID:AB_2631091 |
| Rabbit anti-Ir75c | 1:100 | RRID:AB_2631094 |
| Guinea pig anti-Ir75c | 1:100 | *This study* |
| Guinea pig anti-Ir8a | 1:300 | RRID:AB_2566833 |
| Guinea pig anti-E93 | 1:100 | (30) |
| Rabbit anti-RFP | 1:250 | Abcam ab62341 |
| Chicken anti-GFP | 1:500 | Abcam ab13970 |
| Anti-Chicken Alexa Fluor 488 | 1:500 | Abcam ab150169 |
| Anti-Rabbit Alexa Fluor 488 | 1:100 | RRID:AB_2576217 |
| Anti-Rabbit Cy3 | 1:250 | Milan Analytica AG 111-165-1440 |
| Anti-Rabbit Cy5 | 1:100 | Jackson ImmunoResearch<br>111-175-144 |
| Anti-Guinea pig Cy5 | 1:300 | Abcam ab102372 |
| Anti-Rabbit Alexa Fluor 405 | 1:500 | RRID:AB_221605 |
| Anti-Digoxigenin-POD | 1:300 | Roche Diagnostics 11 207 733 910 |
| Anti-FLUO-POD | 1:300 | Roche Diagnostics 11 426 346 910 |

